# Contralateral Delay Activity is not sensitive to cognitive decline in older adults at risk of Mild Cognitive Impairment

**DOI:** 10.1101/753392

**Authors:** Francesca R Farina, Gabija Pragulbickaitė, Marc Bennett, Cian Judd, Kevin Walsh, Samantha Mitchell, Redmond G O’Connell, Robert Whelan

**Affiliations:** Trinity College Institute of Neuroscience, Trinity College Dublin, Dublin 2, Ireland; MRC- Cognition and Brain Science Unit, University of Cambridge, UK; Global Health Brain Institute, Trinity College Dublin, Dublin 2, Ireland

**Keywords:** Working Memory, Mild Cognitive Impairment, EEG, Older adults, Contralateral Delay Activity

## Abstract

Contralateral delay activity (CDA) has been proposed as a pre-clinical marker for Mild Cognitive Impairment (MCI). However, existing evidence is limited to one study with a small sample size (n=12 per group). Our aim was to compare CDA amplitudes in a larger sample of low- and high-risk older adult groups (n=35 per group). As expected, behavioural performance decreased as the number of memory items increased, and the low-risk group out-performed the high-risk group. However, we found no differences in CDA amplitudes across groups, indicating that WM capacity increased irrespective of risk-level. These findings suggest that the CDA is not a sensitive marker of MCI risk. More broadly, our results highlight the difficulty in identifying at-risk individuals, particularly as MCI is a heterogeneous, unstable condition. Future research should prioritise longitudinal approaches in order to track the progression of the CDA and its association with cognitive decline in later life.

## 1. Introduction

Contralateral delay activity (CDA) is characterised by a negative event-related potential (ERP; time-locked encephalograms) waveform that is sensitive to the number of items maintained in working memory (WM) and persists throughout the retention period (Feldmann-Wustefeld, Vogel, & Awh, 2018). The CDA has become increasingly popular as an index of age-related WM changes (Du, Ji, Chen, Tang, & Han, 2018). However, existing evidence regarding the effects of both age and performance level on CDA amplitudes is equivocal (Jost, Bryck, Vogel, & Mayr, 2011; Sander, Werkle-Bergner, & Lindenberger, 2011; Schwarzkopp, Mayr, & Jost, 2016; Stormer, Li, Heekeren, & Lindenberger, 2013). One possible explanation is that atypical CDA waveforms reflect pathological decline, rather than normal aging. In support of this, Newsome, Pun, Smith, Ferber and Barense (2013) reported CDA differences between healthy older adults and those at-risk for Mild Cognitive Impairment (MCI). At-risk older adults did exhibit the expected differentiation of the CDA across set sizes. In contrast, amplitudes in the healthy older adult group were indistinguishable from those of a younger adult group. Based on their findings, the authors proposed the CDA to be a pre-clinical marker for MCI. Although promising, this study included a small sample size (n=12 per group), and no other studies since have investigated whether the CDA is sensitive to pre-clinical MCI. Here, we aimed to replicate the findings of Newsome et al. by comparing amplitudes in healthy older adults separated into low- and high-risk groups. We hypothesized that older adults at higher risk of MCI would exhibit reduced differentiation of CDA amplitudes across set sizes, relative to low-risk older adults.

## 2. Methods

Seventy older adults were included, grouped based on their performance on the Montreal Cognitive Assessment (MoCA; Nasreddine et al., 2005) into low-risk (≥26; n=35) and high-risk groups (≤25; n=35). Groups were matched for age and education. The School of Psychology, Trinity College Dublin, Research Ethics Committee, provided ethical approval. Research accorded to Declaration of Helsinki principles. Informed consent was obtained prior to testing. Participants completed the CDA task (Vogel & Machizawa, 2004) while undergoing EEG recording. EEG data were recorded using Active Two Biosemi™ system from 134 electrodes (128 scalp) organised according to the 10-5 system (Oostenveld & Praamstra, 2001). CDA amplitude was calculated by subtracting ipsilateral from contralateral activity (300-800ms post-memory array onset), averaged across hemispheres for three posterior electrodes (P5/P6, PO7/PO8 and CP3/CP4), at each set size (1, 2 and 3; correct trials only). Behavioural performance was assessed via mixed factorial ANOVA, with group and set size as factors. CDA amplitudes were also compared using mixed factorial ANOVAs, with group, set size and electrode as factors. Finally, CDA amplitudes were examined using a Bayesian ANOVA to quantify support for the null/alternative hypothesis (Wagenmakers et al., 2018), using group, set size and electrode as factors. Full details of the experimental setup, data processing and statistical analyses can be found in the Supplementary Methods.

## 3. Results

Table 1 displays demographic and neuropsychological information per group (see also Supplementary Results). There were significant main effects of group and set size for task accuracy (*ps* ≤ .03). Low-risk older adults performed better than high-risk older adults and accuracy decreased as set size increased. There were significant main effects of group and set size, and a significant interaction for WM capacity (*ps* ≤ .03). Capacity estimates were higher for the low- vs. high-risk group, and they increased across set sizes. Reaction times increased with set size (*p* < .001). No other main effects or interactions were found (see Supplementary Results). ERP analyses yielded similar results; there were main effects of set size for CDA amplitude (between 300-800 ms). The CDA became larger as set size increased (*ps* ≤ .002). Early in the retention period (300-400ms post-memory array), CDA amplitude differences were found between set size 1 and set sizes 2 (*p* = .05) and 3 (*p* < .001). For the remainder of the time (400-800ms), CDA amplitudes differed across all set sizes (*ps* < 0.01). Crucially, there were no main effects of group or interactions between group, electrode and/or set size (see Figures 1, S1). Estimated Bayes factors supported these results, producing strong evidence in favour of set size differences (BF_10_ > 100) and moderate-to-strong evidence against differences between groups (BF_10_ = .34) or electrodes (BF_10_ = .05). Two-way and three-way interaction effects also received weak support (all BF_inclusion_ ≤ .08).

**Table 1:** Demographic information, neuropsychological and behavioural performance for low- and high-risk groups.

|  | Low Risk |  |  | High Risk |  |  |
| --- | --- | --- | --- | --- | --- | --- |
| Total N | 35 |  |  | 35 |  |  |
| Age (years) | 66.37 (5.81) |  |  | 67.80 (7.75) |  |  |
| Sex (female) | 25 (71.4) |  |  | 17 (48.6) |  |  |
| Edu (years) | 14.33 (2.81) |  |  | 13.12 (3.32) |  |  |
| MoCA | 27.60 (1.29) |  |  | 23.14 (2.13) |  |  |
| MMSE | 28.91 (1.15) |  |  | 27.89 (2.65) |  |  |
| Handedness | 34 right-handed |  |  | 32 right-handed† |  |  |
| NART FSIQ | 118.24 (5.22) |  |  | 113.31 (7.38)† |  |  |
| WMS-IV | 21.26 (6.00) |  |  | 15.12 (5.70) |  |  |
|  | **Set size 1** | **Set size 2** | **Set size 3** | **Set size 1** | **Set size 2** | **Set size 3** |
| WM Capacity | 0.96 (.01) | 1.86 (.02) | 2.13 (.07) | 0.87 (.03) | 1.68 (.06) | 1.89 (.06) |
| CDA ACC (%) | 97.83<br>(1.67) | 93.98<br>(3.48) | 85.52<br>(6.84) | 93.59<br>(11.16) | 89.38<br>(8.93) | 79.74<br>(9.81) |
| CDA RTs (ms) | 1.04 (.52) | 1.27 (.52) | 1.44 (.42) | 0.99 (.37) | 1.2 (.38) | 1.38 (.42) |

*Note:* Values indicate mean scores with standard deviation (SD) in brackets. Abbreviations: ACC: accuracy; CDA: Contralateral Delay Activity; Edu: Education; FSIQ: Full-scale IQ; MoCA: Montreal Cognitive Assessment; MMSE: Mini-Mental State Examination; NART: National Adult Reading Test; RTs: reaction times; WM: working memory; WMS-IV: Wechsler Memory Scale 4<sup>th</sup> Edition. †A small number of scores were missing: 1 for handedness, 2 for NART and 1 for WMS.

**Figure 1:**
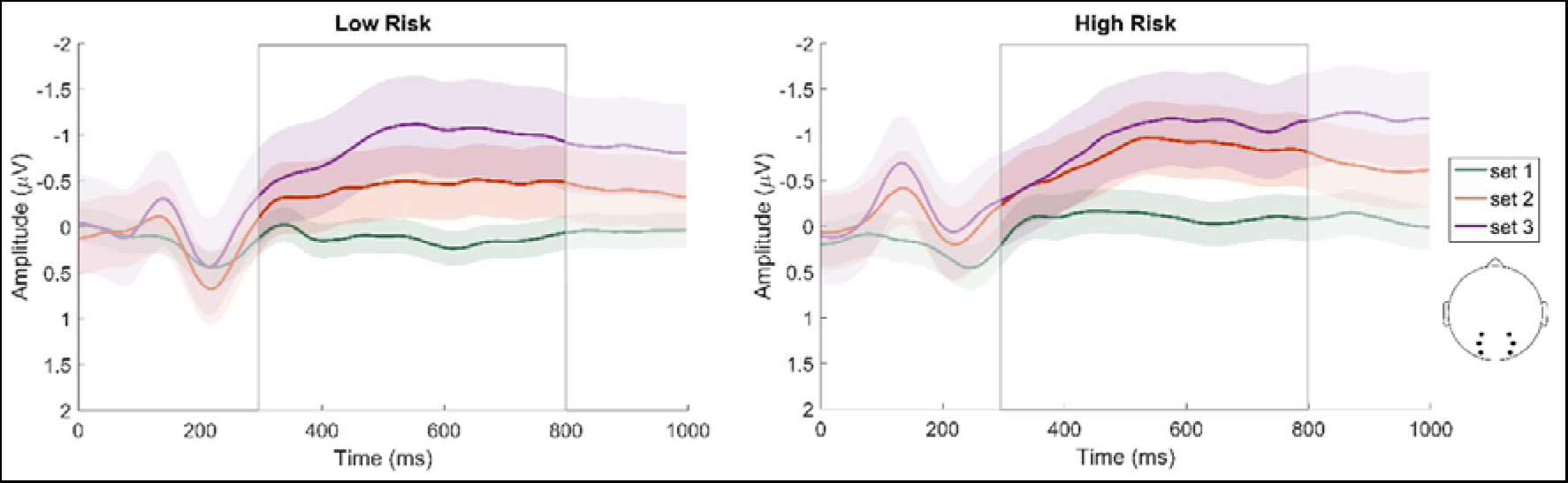
Contralateral delay activity (CDA) for low- and high-risk groups for set sizes 1, 2 and 3. Activity averaged from six electrodes (CP3/CP4, P5/P6 and PO7/PO8) was analysed from 300-800ms (grey box outline). Coloured shaded areas indicate standard deviation.

## 4. General Discussion

We aimed to investigate CDA sensitivity to MCI risk. As expected, behavioural performance decreased as the number of memory items increased, and the low-risk group out-performed the high-risk group. However, although CDA amplitudes also increased with set size, we found no ERP group differences. Both groups exhibited equivalent differentiation of CDA amplitudes, indicating that WM capacity increased irrespective of risk-level. These findings suggest that the CDA is not a sensitive marker for pre-clinical MCI, contrary to Newsome et al. (2013). It is possible that sample size contributed to the discrepancy in findings (i.e. n=25 previously vs n=70 here) (Button et al., 2013; Larson & Carbine, 2017). For example, Button et al. (2013) discuss the ‘Winner’s curse’, in which a lower powered study first reports a significant result, but subsequent studies report a smaller effect size. It is certainly possible, therefore, that the CDA is a marker of MCI risk although the effect size may be too small to be of clinical relevance. We note a methodological difference that may have influenced our results: our task had set sizes of 1, 2 and 3, vs. 1, 3 and 4 in Newsome et al. However, previous results showed no differentiation across any set size in high-risk adults, whereas we found differences across all set sizes. More broadly, our results highlight the difficulty in identifying at-risk individuals, particularly as MCI is a heterogeneous, unstable condition (Petersen et al., 2014). Future research in this area should prioritise longitudinal approaches in order to track the progression of the CDA and its association with cognitive decline in later life.

## Supporting information

Supplemental Methods & Results

## Acknowledgements

The data collection was partially supported by DART Neuroscience. The company was not involved in the conduct of the research or preparation of the article. This work is supported by Irish Research Council’s Government of Ireland Postdoctoral Fellowship grants to Francesca Farina (EPSPD/2017/110) and Marc Bennett (GOIPD/2016/617). Robert Whelan is supported by Science Foundation Ireland (project number: 16/ERCD/3797).

## References

Button, K. S., Ioannidis, J. P., Mokrysz, C., Nosek, B. A., Flint, J., Robinson, E. S., & Munafo, M. R. (2013). Power failure: why small sample size undermines the reliability of neuroscience. Nat Rev Neurosci, 14(5), 365–376. doi:10.1038/nrn3475

Delorme, A., & Makeig, S. (2004). EEGLAB: an open source toolbox for analysis of single-trial EEG dynamics including independent component analysis. J Neurosci Methods, 134(1), 9–21. doi:10.1016/j.jneumeth.2003.10.009

Du, X., Ji, Y., Chen, T., Tang, Y., & Han, B. (2018). Can working memory capacity be expanded by boosting working memory updating efficiency in older adults? Psychol Aging, 33(8), 1134–1151. doi:10.1037/pag0000311

Folstein, M. F., Folstein, S. E., & McHugh, P. R. (1975). “Mini-mental state”. A practical method for grading the cognitive state of patients for the clinician. J Psychiatr Res, 12(3), 189–198.

Jost, K., Bryck, R. L., Vogel, E. K., & Mayr, U. (2011). Are old adults just like low working memory young adults? Filtering efficiency and age differences in visual working memory. Cereb Cortex, 21(5), 1147–1154. doi:10.1093/cercor/bhq185

Larson, M. J., & Carbine, K. A. (2017). Sample size calculations in human electrophysiology (EEG and ERP) studies: A systematic review and recommendations for increased rigor. Int J Psychophysiol, 111, 33–41. doi:10.1016/j.ijpsycho.2016.06.015

Nasreddine, Z. S., Phillips, N. A., Bedirian, V., Charbonneau, S., Whitehead, V., Collin, I., Chertkow, H. (2005). The Montreal Cognitive Assessment, MoCA: a brief screening tool for mild cognitive impairment. J Am Geriatr Soc, 53(4), 695–699. doi:10.1111/j.1532-5415.2005.53221.x

Newsome, R. N., Pun, C., Smith, V. M., Ferber, S., & Barense, M. D. (2013). Neural correlates of cognitive decline in older adults at-risk for developing MCI: Evidence from the CDA and P300. Cognitive Neuroscience, 4(3-4), 152–162. doi:10.1080/17588928.2013.853658

Oostenveld, R., & Praamstra, P. (2001). The five percent electrode system for high-resolution EEG and ERP measurements. Clin Neurophysiol, 112(4), 713–719.

Petersen, R. C., Caracciolo, B., Brayne, C., Gauthier, S., Jelic, V., & Fratiglioni, L. (2014). Mild cognitive impairment: a concept in evolution. Journal of internal medicine, 275(3), 214–228. doi:10.1111/joim.12190

Sander, M. C., Werkle-Bergner, M., & Lindenberger, U. (2011). Contralateral delay activity reveals life-span age differences in top-down modulation of working memory contents. Cereb Cortex, 21(12), 2809–2819. doi:10.1093/cercor/bhr076

Schwarzkopp, T., Mayr, U., & Jost, K. (2016). Early selection versus late correction: Age-related differences in controlling working memory contents. Psychol Aging, 31(5), 430–441. doi:10.1037/pag0000103

Stormer, V. S., Li, S. C., Heekeren, H. R., & Lindenberger, U. (2013). Normative shifts of cortical mechanisms of encoding contribute to adult age differences in visual-spatial working memory. Neuroimage, 73, 167–175. doi:10.1016/j.neuroimage.2013.02.004

Vogel, E. K., & Machizawa, M. G. (2004). Neural activity predicts individual differences in visual working memory capacity. Nature, 428(6984), 748–751. doi:10.1038/nature02447

Wagenmakers, E.-J., Love, J., Marsman, M., Jamil, T., Ly, A., Verhagen, J., Morey, R. D. (2018). Bayesian inference for psychology. Part II: Example applications with JASP. Psychonomic Bulletin & Review, 25(1), 58–76. doi:10.3758/s13423-017-1323-7

