## Supplemental Methods & Results for "Contralateral Delay Activity is not sensitive to cognitive decline in older adults at risk of Mild Cognitive Impairment"

**Supplementary Materials**

**Supplementary Methods**

*Participants*

Seventy older adults (Age: M= 67.07, SD = 6.96; 42 females; Education: M = 13.73, SD = 3.11) recruited from the general population were included in the study. Inclusion criteria were: being 55 years or older, with no history of neurological (epilepsy, multiple sclerosis, depression, Parkinson’s disease, dementia or brain trauma) or psychiatric (anxiety, depression, bipolar) disorders; and no history or current use of psychiatric medication or substance abuse. All participants had normal or corrected-to-normal vision and scored within the normal range on the Mini-Mental State examination (MMSE) (>24; Folstein, Folstein, & McHugh, 1975). Participants were divided into low- and high-risk groups based on their scores on the MoCA. Participants who scored 26 or above were classed as low risk; those who scored 25 or below were classed as high risk. Groups were matched for age and education. Ethical approval was obtained from the School of Psychology Research Ethics Committee, Trinity College Dublin. All research was conducted according to the principles expressed in the Declaration of Helsinki. Written informed consent was obtained from participants prior to testing. Participants were paid €60 plus travel expenses.

*Neuropsychological Examination*

Cognitive status was measured using the MMSE (Folstein, Folstein & McHugh, 1975) and MoCA (Nasreddine et al., 2005). Intelligence was measured using the National Adult Reading Test (NART; Nelson, 1982).

*Experimental setup and task*

Participants were seated in a dark, soundproofed room in front of a computer screen. All participants completed CDA task, which has been described previously (Vogel & Machizawa, 2004). The task consisted of three blocks of 60 trials (360 trials total), lasting approximately 20 minutes. Each trial began with an arrow above a centrally placed fixation cross. Participants focused their attention in the direction of the arrow while maintaining central fixation. Following a 1000ms delay, a bilateral memory array consisting of coloured squares appeared for 150ms, followed by a test array, which remained on the screen until participants responded with a button press. Participants were required to indicate if the squares remained the same or changed colour. This was followed by a 750ms delay before the next trial began.

*EEG data acquisition and pre-processing*

Participants underwent EEG recording at the same time as completing the CDA task. EEG data were recorded using Active Two Biosemi™ system from 134 electrodes (128 scalp) organised according to the 10-5 system (Oostenveld & Praamstra, 2001). The vertical and horizontal electro-oculograms were recorded bilaterally, approximately 3cm below the eye and from the outer canthi, respectively. EEG data was pre-processed using the EEGLAB toolbox (Delorme & Makeig, 2004) with the FASTER plug-in (Nolan, Whelan, & Reilly, 2010). The data were bandpass filtered between 1 and 95 Hz, notch filtered at 50Hz and average referenced across all scalp electrodes. FASTER identified and removed non-neural independent components, epochs with large artifacts (e.g. muscle movements), and channels with poor signal quality (interpolated). After pre-processing, data were visually inspected for any remaining noise. Independent components (ICs) and epochs containing artifacts were removed using ManualQC. Following pre-processing, EEG data were segmented into epochs from -200ms to 2000ms around the memory array. Epochs were average referenced and baseline-corrected by subtracting the mean amplitude of the 100ms pre-stimulus period. Epochs were extracted for set sizes 1, 2 and 3, presented left and right of fixation, respectively.

*CDA calculation*

CDA amplitude was calculated by subtracting ipsilateral from contralateral activity from 350-850ms following the onset of the memory array, averaged across hemispheres for three posterior electrodes (P5/P6, PO7/PO8 and CP3/CP4). Electrode selection was based on Newsome *et al.* (2013). CDA amplitude was assessed for each participant in each set size (1, 2 and 3) for correct trials only.

*Data analyses*

Statistical analyses were performed using SPPS Version 24. Group differences in demographic information and neuropsychological test performance were examined using independent samples t-tests. Behavioural performance on the CDA task was assessed via mixed factorial ANOVA, with group (low, high) as the between-groups factor and set size (1, 2, 3) as the within-groups factor. Three ANOVAs were run, assessing mean accuracy, WM capacity (*K*) and reaction times (RTs), respectively. CDA amplitudes (from 300-800ms) were also compared across groups using mixed factorial ANOVAs, with group (low, high) as the between-groups factor and set size (1, 2, 3) and electrode (P5/P6, PO7/PO8 and CP3/CP4) as within-groups factors. To investigate the potential time course of any CDA effects, CDA amplitude was divided into five 100ms intervals (between 300-800ms) and analysed using separate mixed factorial ANOVAs. Bonferroni post hoc tests were carried out where appropriate. Where Mauchly’s Test of Sphericity was violated, values reported are Greenhouse-Geisser corrected. A priori power analyses indicated that we needed to have a minimum sample size of 66 participants to have 95% power for detecting a medium effect size at the traditional .05 criterion of statistical significance. Mean CDA amplitudes were also examined using a Bayesian mixed factorial ANOVA to quantify the evidence for the null and alternative hypotheses (Wagenmakers et al., 2018). Group (low, high) was the between-groups factor and set size (1, 2, 3) and electrode (P5/P6, PO7/PO8 and CP3/CP4) were within-groups factors. Cauchy post hoc tests were carried out where appropriate.

**Supplementary Results**

*Demographic and neuropsychological information*

Groups did not differ in age (*t*(63) = .87, *p = .*39) or years of education (*t*(65) = - 1.64, *p =.*11). However, there were significant groups differences in MMSE scores (*t*(46) = - 2.10, *p =.*04), NART performance (*t*(66) = - 2.99, *p =.*004) and the Symbol Span task (*t*(67) = - 4.40, *p < .*001).

*Behavioural data: CDA task*

For accuracy, main effects of group (*F*(1,68) = 9.21, p = .003, ƞp² = .12) and set size (*F*(1,101) = 139.02, *p* < .001, ƞp² = .67) were found (see Table 1). The low-risk group performed better (M = 92.44%, SD = 1.14) than the high-risk group (M = 87.57%, SD = 1.14). Accuracy also decreased across set sizes (set 1: M = 95.71, SD = .96, set 2: M = 91.68, SD = .81, set 3: 82.63, SD = 1.01). There was no significant group x set size interaction effect (*F*(2,101) = .51, p = .55, ƞp² = .007). For WM capacity, there was a significant interaction between group and set size (*F*(1,82) = 4.80, *p* = .03, ƞp² = .07), a main effect of group (*F*(1,68) = 9.56, *p* = .003, ƞp² = .12) and of set size (*F*(1,82) = 329.68, *p* < 0.001, ƞp² = .83). Capacity estimates for the high-risk group differed significantly between set size 1 and 2 (*p* <.001), 2 and 3 (*p* = 0.1), 3 and 1 (*p* < .001) (set 1: M = .87, SD = .03, set 2: M = 1.68, SD = .06, set 3: M = 1.89, SD = .06) Capacity estimates for the low-risk group also differed significantly at each set size (all *p*s < .001) (set 1: M = .96, SD = .01; set 2: M = 1.86, SD = .02, set 3: M = 2.13, SD = .07). In addition, group differences in capacity estimates were noted at each set size (set size 1: *F*(1,69) = 5.06, *p* = .03, set size 2: *F*(1,69) = 8.14, *p* = .006, set size 3: *F*(1,69) = 8.01, *p* = .006). In all cases, estimates were higher for the low-risk group. There was also a main effect of set size for reaction times (RTs), (*F*(2,136) = 58.61, *p* < .001, ƞp² = .46), whereby RTs increased across set sizes (*p*s < .001). No main effect of group (*F*(1,68) = .48, *p* = .49, ƞp² = .01) or group x set size interaction was noted (*F*(2,136) = 1.34, *p* = .27, ƞp² = .01).

*EEG data: CDA task*

On average, 10% (SD = 7.28) of epochs were removed. Analysis of mean CDA amplitudes (between 300-800ms) revealed a significant main effect of size (*F*(2,136) = 34.84, *p* < .001, ƞp² = .34). The CDA became larger (i.e. more negative) as the number of to-be-remembered items increased (*p*s ≤ .002; see Figure S1). No main effects of group (*F*(1,68) = 1.43, *p* = .24, ƞp² = .02) or electrode were noted (*F*(2,136) = 1.34, *p* = .27, ƞp² = .02). Two-way interactions between group and set size (*F*(2,136) = .89, *p* = .42, ƞp² = .01), set size and electrode (*F*(4,272) = .34, *p* = .85, ƞp² = .01) or group and electrode were found (*F*(2,136) = .59, *p* = .56, ƞp² = .01). The three-way interaction between group, set size and electrode was also non-significant (*F*(4,272) = 1.29, *p* = .27, ƞp² = .02).

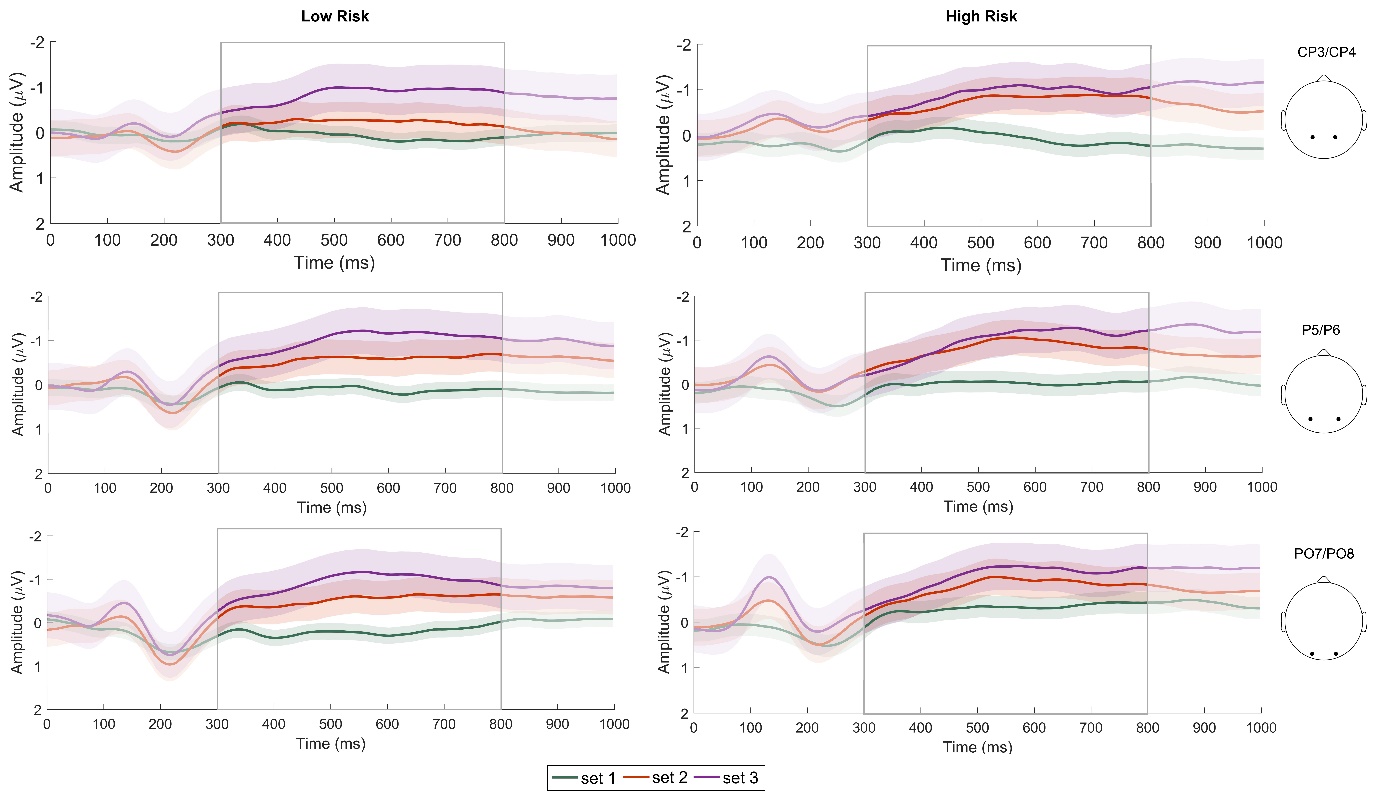

**Figure S1:** Contralateral delay activity (CDA) for low- and high-risk groups for set sizes 1, 2 and 3 from three electrode pairs (CP3/CP4, P5/P6 and PO7/PO8) between 300-800ms (grey box outline). Coloured shaded areas indicate standard deviation.

Given the absence of electrode effects, we collapsed across electrode site for subsequent analyses. Mean CDA amplitude was divided into 100ms intervals (between 300-800ms) to investigate any potential time-specific effects. Between 300-400ms, the main effect of set size was significant (*F*(2,136) = 8.75, *p* < .001, ƞp² = .11) but the main effect of group (*F*(1,68) = .17, *p* = .69, ƞp² < .001) and group x set size interaction were not (*F*(2,136) = .59, *p* = .56, ƞp² = .01). CDA amplitude at set size 1 was significantly reduced compared to set sizes 2 (*p* = .05) and 3 (*p* < .001). A main effect of set size was also found between 400-500ms (*F*(2,136) = 28.00, *p* < .001, ƞp² = .29), but the effect of group (*F*(1,68) = 1.22, *p* = .27, ƞp² = .02) and interaction effect were not significant (*F*(2,136) = .62, *p* = .54, ƞp² = .01). CDA amplitude increased across all set sizes (*p*s < 0.01). Identical results were obtained for 500-600ms (set size: *F*(2,136) = 40.54, *p* < .001, ƞp² = .37; group: *F*(1,68) = 2.12, *p* = .15, ƞp² = .03; group x set size interaction: *F*(2,136) = 1.53, *p* = .22, ƞp² = .02; set size pairwise comparisons: *p*s < .001), 600-700ms (set size: *F*(2,136) = 42,13, *p* < .001, ƞp² = .38; group: *F*(1,68) = 2.02, *p* = .16, ƞp² = .03; group x set size interaction: *F*(2,136) = .63, *p* = .54, ƞp² = .01; set size pairwise comparisons: *p*s ≤ .001) and 700-800ms (set size: *F*(2,136) = 26.80, *p* < .001, ƞp² = .28; group: *F*(1,68) = 1.52, *p* = .22, ƞp² = .02; group x set size interaction: *F*(2,136) = .51, *p* = .60, ƞp² = .01; set size pairwise comparisons: *p*s < .01).

Finally, we conducted a Bayesian mixed factorial ANOVA to quantify support for the abovementioned effects. An estimated Bayes factor produced overwhelming evidence in favour of set size effects (BF_01_ < .001). Support for the alternative hypothesis (i.e. that there was difference in CDA amplitude) was indicated for all set size comparisons (Posterior Odds values ≤ .01). In contrast, strong evidence against electrode effects was found (BF_01_ = 18.74). An estimated Bayes factor for group also yielded moderate evidence against group effects (BF_01_ = 2.91). Two-way interactions between group and set size (BF_inclusion_ = .08), group and electrode (BF_inclusion_ < .01) and set size and electrode (BF_inclusion_ = .001) received weak support, as did the three-way group x set size x electrode interaction (BF_inclusion_ < .001).
